# Highly resolved spatial transcriptomics for detection of rare events in cells

**DOI:** 10.1101/2021.10.11.463936

**Authors:** Silvia Groiss, Daniela Pabst, Cynthia Faber, Andreas Meier, Annette Bogdoll, Conny Unger, Benedikt Nilges, Sascha Strauss, Esther Föderl-Höbenreich, Melina Hardt, Andreas Geipel, Frank Reinecke, Christian Korfhage, Kurt Zatloukal

## Abstract

Single-cell spatial transcriptomics technologies leveraged the potential to transcriptionally landscape sophisticated reactions in cells. Current methods to delineate such complex interplay lack the flexibility in rapid target adaptation and are particularly restricted in detecting rare transcripts. We developed a multiplex single-cell RNA *In-situ* hybridization technique, called ‘Molecular Cartography’ (MC) that can be easily tailored to specific applications and, by providing unprecedented sensitivity, specificity and resolution, is particularly suitable in tracing rare events at a subcellular level. Using a SARS-CoV-2 infection model, MC allows the discernment of single events in host-pathogen interactions, dissects primary from secondary responses, and illustrates differences in antiviral signaling pathways affected by SARS-CoV-2, simultaneously in various cell types.

## Introduction

Single-molecule RNA-fluorescence in situ hybridization (smFISH) together with single-cell RNA sequencing (scRNA-seq) techniques are the current pillars to record transcriptional responses to a disease or environmental agent and have proved highly useful in investigating e.g., Severe Acute Respiratory Syndrome Coronavirus 2 (SARS-CoV-2), the agent responsible for the recent pandemic^1-3^. Several advanced multiplexed smFISH-based applications (MERFISH, Visium, Cartana, ZipSeq, SeqFISH, DSP) are available that provide analysis of spatio-transcriptional activities^4-9^. These technologies are crucial assets to better understand the functional heterogeneity of cells as well as cell autonomous and paracrine effects as inherent principles of physiological and disease conditions, and to correlate such indices to genetic and developmental differences^7,8,10^. However, they are often limited in the number of simultaneously tested samples, time-consuming in adapting protocols to new targets or limited in the resolution or sensitivity necessary to spatially map the transcriptional landscape at a subcellular level, particularly when tracing and dissecting rare transcriptional events^7,8,11-13^.

In particular, separating timely or spatially close but disparate events, e.g. responses in primary and secondary infection events, remains highly challenging. Yet, understanding the response dynamics between such primary and secondary infected cells especially in comparison to non-infected bystander cells is crucial to capture the priming of bystander cells towards a collective antiviral response. Shedding light on this very process may help us to better comprehend the disparate infectivity rates and discordant inflammatory signaling observed in COVID-19 patients and its underlying pathogenesis on a molecular level^14^.

To bridge this gap, we developed ‘Molecular Cartography’ (MC), a highly resolved multiplexed smFISH technique that, through enhanced sensitivity and specificity, allows tracing of even scarcely expressed, single RNA molecules of up to 100 genes simultaneously at a subcellular resolution in multiple samples in parallel, or the analysis of up to 2400 genes in a single experiment. Using a set of fluorescently labeled probes that can easily be designed for specific applications, MC detects and codes single transcripts from individual genes in consecutive rounds that are decoded and counted by a specifically devised algorithm. The rapid design and flexibility to change these customizable probe sets, in particular, unlocks the analytical potential to dynamically adapt research hypotheses within a few days.

Apart from illustrating the advances in specificity, sensitivity and reproducibility, we demonstrate its feasibility by investigating heterogeneous gene expression patterns of key viral entry factors within clonal populations in lung, colorectal, and hepatocellular cell lines and compare primary and secondary pathogen host responses in an infection model of SARS-CoV-2^15^. We further investigate differences in overall infectivity and track pathways known to be affected by coronaviridae^16-18^. Finally, by simultaneously detecting and localizing context-enriched targets, we directly trace the response trajectory starting at primary infection sites (INFS) and highlight the complex propagation network of SARS-CoV-2^19^.

## Results

MC harbors the advantages of previous multiplexed smFISH based technologies but with increased flexibility (i.e. rapid adaptation to specific research questions), sensitivity and specificity compared to current applications including concurrent analysis of multiple samples. The previously designed and synthesized probes are hybridized to their target species (**Fig. 1a**) followed by repeated rounds of probe colorization with fluorophores specific for a certain target species. Transcript-specific probes are designed using an algorithm developed specifically for MC by Resolve BioSciences that allows the design of probe sets of almost all genes of an organism. *In-silico* probe design is one of the most critical steps for spatial transcriptomics analysis. In general, a proper probe design influences uniformity, sensitivity and specificity which has been shown in previous applications targeting probe design e.g., for microarrays^20^. Probe design for accurate sensitivity and specificity was optimized in several iterations of MC experiments including the design algorithm, which ultimately allowed the reduction of the probe number per transcript. For example, MC effectively quantifies short transcripts such as the human *CCL8* transcripts (length ∼850 nucleotides) or mouse interleukin (IL) 18 (length: ∼700 nucleotides) in individual cells. This indicates that MC detects and quantifies even short transcripts or transcript variants.

**Fig. 1.**
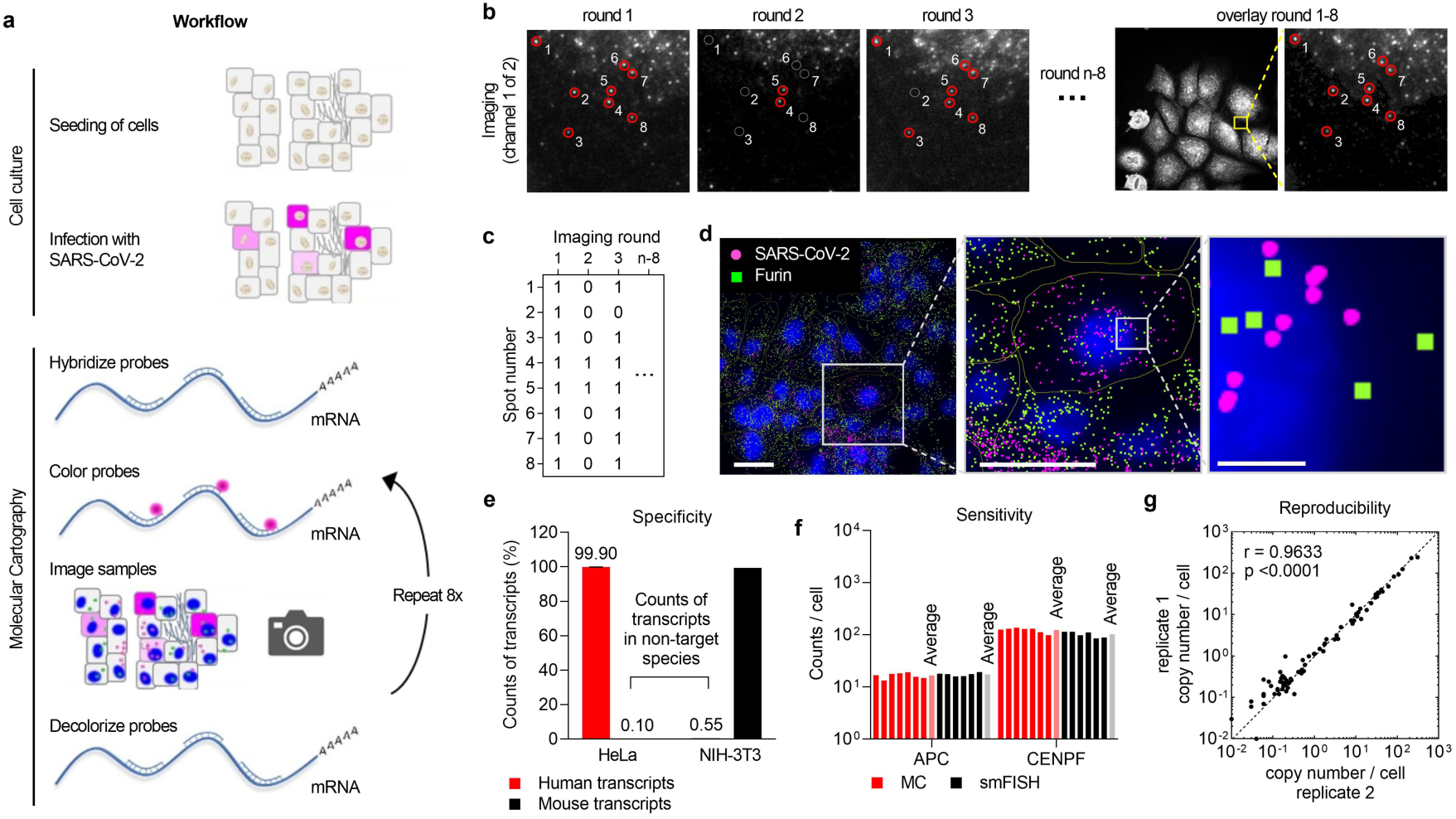
Workflow, data processing and validation of MC. **a**, Workflow of MC. Cartoon illustrating how SARS-CoV-2 infected and fixed cells are hybridized with probes, color coded, imaged and decolorized for repeated rounds of coloring and imaging. **b**, Image processing using the devised algorithm. Left panel, representative images of round 1-3. Right panel, overlay of round 1-8. **c**, Corresponding code for the spots identified in b. **d**, Illustration of two out of 81 transcripts (SARS-CoV-2 and *FURIN*) measured in Huh7 cells. Left panel, overview of the region of interest (ROI). Middle panel, single cell close-up view of a single cell within the ROI. Right panel, targets detected at a subcellular resolution close to the nucleus. Left and middle scale bar, 50 μm. Right scale bar, 5 μm. **e**, Specificity measurement of MC. Both human and mouse transcripts were hybridized to both human HeLa and mouse NIH-3T3 cells and detected using MC (Supplementary Fig. 1). Data is shown as mean ± standard deviation (SD) from n = 3 independent technical replicates. **f**, Sensitivity measurement of MC using an orthogonal approach. Two probe sets labeled by two different fluorophores were hybridized to HeLa cells and tested using MC in comparison to standard smFISH. The copy number per target was calculated as the average count per cell per experiment. Results derived from MC and smFISH were comparable across all experiments (n = 6 (smFISH) – 7 (MC) total technical replicates). **g**, Correlation of the average count per RNA species per cell of two independent MC experiments using Huh7 cells (n = 2 biological replicates analyzing 212 and 290 single cells, respectively). The Spearman correlation coefficient is 0.96 with a p-value of <0.0001. The x = y axis is marked by a dashed line.

In total, eight consecutive rounds of colorization, imaging, de-colorization and re-colorization build up a combinatorial color code that is specific for every target sequence. All combinatorial coding schemes have in common that the number of combinations increases with the length of a single code N. The length of the code is given by the number of detection rounds and number of fluorophores (x) used for the multiplex smFISH approach (x^N^). Although the number of combinations increases with each round, we reduced the number of potential available codes therefore increasing the distance between the codes. This increases the Hamming distance, which in turn, results in several advantages. The large distance between codes allows for correction of the code, if a certain signal is not detected or wrongly assigned. An increase in code distances thus strongly improves specificity and sensitivity by reducing rates of misidentification due to error recognition and correction.

An example of three consecutive rounds of imaging (channel 1 of 2) is given in **Fig. 1b**. The first three images demonstrate how a color code is built up in three different rounds. Note that some fluorescent spots are detectable in all three rounds of the same channel, while others appear only in round 2 and/or 3, respectively. After de-colorization, individual spots disappear and re-appear depending on the code in the next colorization round, ultimately generating a unique molecular identifier per target (**Fig. 1c**). Although colorization and de-colorization is stable over more than ten rounds, we used only eight rounds for a faster analysis and to build up the code to obtain the most sensitive results. A software specifically developed for MC analyzes the images of all eight rounds to compute the color code and map the corresponding target transcript in xyz position within the sample. To facilitate high error robustness, the Hamming distance of the codes ensures the highest specificity of analyses (see **Online Methods**).

Together, this process delineates the spatial distribution of numerous target transcripts at subcellular resolution (**Fig. 1d**, close-up images in the middle and right panel; **Supplementary Video 1**,). Only two (SARS-CoV-2 Np and *FURIN*) of more than 80 targets are highlighted for better visualization. Infection by SARS-CoV-2 is indicated by detecting the number of nucleocapsid protein (Np) transcripts (magenta color).

### Specificity, sensitivity and reproducibility

Misidentification of RNA species is a major drawback in spatial transcriptomics. We therefore tested the specificity i.e. misidentification rate of MC including the entire colorization, imaging and decoding process by applying probe sets designed for human and mouse RNA transcripts to both human Hela and mouse NIH-3T3 cells and evaluated the rate of positive signals in both species. In total, we identified 860,000 human and 289,000 mouse transcripts in human and mouse cells, respectively (**Supplementary Fig. 1**). Overall, the fraction of spots decoded within the wrong cell type is 0.1 % (mouse codes in HeLa cells, **Fig. 1e**) and 0.55 % (human codes in mouse cells) and is slightly higher than the rate of false positives (hits in codes not in use) across three independent experiments. This experiment showed that the total MC process from probe design to hybridization, coding, and decoding showed an average specificity of 99.45 % to 99.9 %.

Additionally, transcriptomics approaches are subject to a drop in calling rate due to incomplete detection of target RNAs. Hence, to demonstrate its sensitivity, we compared MC with a single-round smFISH approach. To differentiate true positives and false positives detected by single-round smFISH, we used two different probe sets labeled by two different fluorophores for a certain transcript. Both probe sets bind in an alternating mode on the transcript to exclude a bias by alternative splicing or hairpin structure formation. In comparative experiments, we performed MC on several instruments and by different users. We tested the copy numbers of the transcripts *APC* and *CENPF*, and calculated the average counts per cell in each experiment. Both MC and smFISH resulted in similar sensitivities with regard to both genes (**Fig. 1f**). This indicates that MC preserves the sensitivity of a single smFISH experiment across multiple rounds of signal detection.

To further test for technical bias in MC experiments, we correlated the measured counts per RNA species per cell derived from two independent biological replicates in SARS-CoV-2 infected Huh7 cells and found an excellent correlation as determined by nonparametric Spearman correlation analysis (p<0.0001; **Fig. 1g**).

Together, the MC multiplex smFISH approach is highly suitable for detection, quantitation and localization of in particular rare transcripts in cells.

### Landscaping heterogeneous expression patterns

Given that clonal populations of cell lines exhibit severe transcriptional heterogeneity that is lost in bulk analysis, we used MC to investigate the expression patterns of multiple RNA species in individual cells^15,21^. We used SARS-CoV-2 infected Huh7 cells to target both highly and lowly expressed genes and investigated their preferential subcellular localization to either cytoplasm or nucleus. We observed high expression rates of genes associated with immunity, and in general antiviral signaling in the context of COVID-19 (for example *JAK2, HIF1A* and *NFκB1*, **Fig. 2a**)^22-24^. These highly abundant targets appeared rather homogeneously distributed and were primarily located in the cytoplasm, probably indicating ongoing translation. In contrast, lowly expressed targets such as *PDGFRA, FOSL1* or *STAT5B* showed strong heterogeneity in both expression level and localization (**Fig. 2a**). PDGFs and FOSL1 regulate cell proliferation and differentiation^25,26^, while the universal transcription factor *STAT5B* exhibits numerous biological roles including proliferation and autoimmunity^27^. These observations were confirmed by single cell analysis of n = 16 randomly chosen cells within the same population (**Fig. 2b**). Furthermore, distribution analysis of various targets revealed the cytoplasm as the primary localization site for the majority of probed targets, with the gene encoding the antiviral response receptor RIG-I (*DDX58)* and the interferon stimulated gene (ISG) *C19orf66* more evenly distributed between the two localizations than for example transcripts of the STAT protein family (**Fig 2c**).

**Fig. 2.**
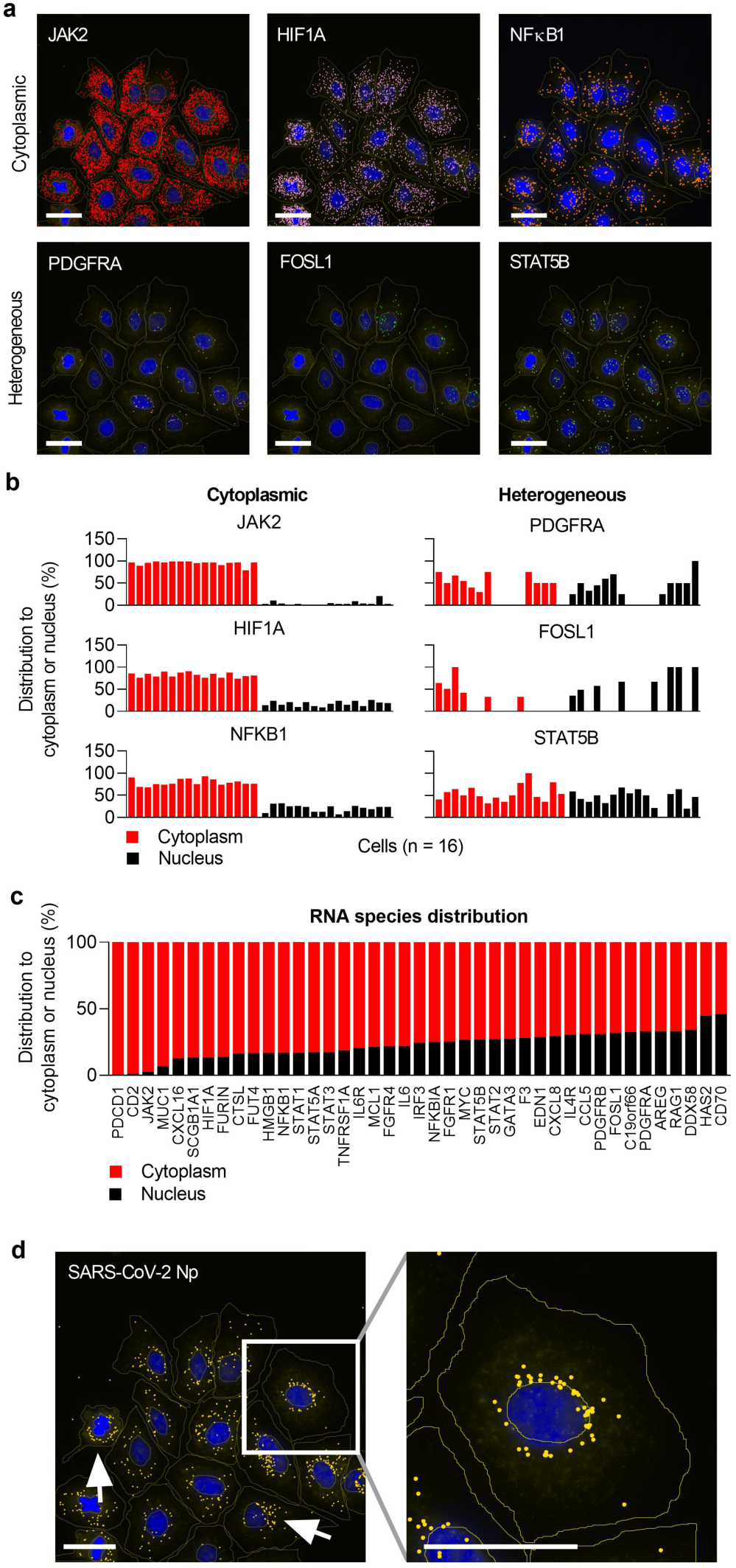
Heterogeneous subcellular distribution of RNA species identified in the MC experiment. **a**, Examples of both the heterogeneous expression and distribution of six distinct RNA species observed across several Huh7 cells. Scale bars, 50 μm. **b**, RNA species shown in a. were found to be either localized in the cytoplasm (*JAK2, HIF1A* and *NFkB1A*) or heterogeneously distributed between cytoplasm and nucleus (*PDGFRA, FOSL1* and *STAT5B*). Data shown of n = 16 individual cells. **c**, Distribution of multiple targets between the cytoplasm and the nucleus averaged within one experiment (n = 18 cells). **d**, Distribution of the SARS-CoV-2 nucleocapsid protein (Np) RNA species within infected Huh7 cells was found to be primarily located close to the nucleus indicating processing in the endoplasmic reticulum (ER). Furthermore, SARS-CoV-2 Np particles appeared polar to one side of the nucleus (arrows and white box with close-up view). Scale bars, 50 μm.

The presented data is derived from 2D images, which could result in misidentification of cytoplasmic targets as nuclei-located when positioned in the cytoplasmic space above the nucleus. However, since highly abundant RNA species, such as *JAK2*, were found to be almost exclusively located in the cytoplasm, we can exclude localization errors due to stochastic distribution of targets and conclude that the localization of target RNAs to different cellular compartments by MC is feasible.

Intriguingly, SARS-CoV-2 Np transcripts were primarily located in the cytoplasm yet close to the nucleus in a site-directed open or closed ring-conformation that suggests an association with the rough endoplasmic reticulum (rER) potentially indicating ongoing translation of Np particles in preparation for exocytosis of SARS-CoV-2 virions^28^ (**Fig. 2d**, arrows, close-up image). These findings highlight the opportunities of using highly multiplexed spatial analysis for investigations in infectivity studies to capture processing of viral transcripts at their subcellular location.

### Tracing antiviral responses in primary infected and bystander cells

We demonstrate the applicability of MC in an infection model of SARS-CoV-2 by using four different cell lines of pulmonary (Calu3), colorectal (Caco2) and hepatocellular origin (Huh7 and PLC5), imaged and analyzed simultaneously in one experiment to allow for meaningful side-by-side comparison. We initially determined the multiplicity of infection (MOI) in VeroE6 cells and finally used an MOI of 0.4 that yielded infection rates at various stages in all four human cell lines that were suitable for studies on antiviral responses in MC experiments.

SARS-CoV-2 Np particle counts decreased with increasing distance to the primary infected cell in Huh7 cells (**Fig. 3a**), which resembled the infection pattern in Calu3, Caco2 and PLC5 cells (**Fig. 3b**). Here, RNAs coding for the key viral entry factors *ACE2, FURIN* and *TMPRSS2* are highlighted^29,30^. To assess a receptor-dependent susceptibility to SARS-CoV-2 infections including a potential associated virus-induced downregulation, we tested their expression patterns in infected and mock-infected cells (n = 13 individual cells per cell line) in relation to the SARS-CoV-2 Np particle count (**Fig. 3c**). We found only low expression levels of *ACE2* across all tested cell lines despite active viral replication, supporting previous findings^14^.

**Fig. 3.**
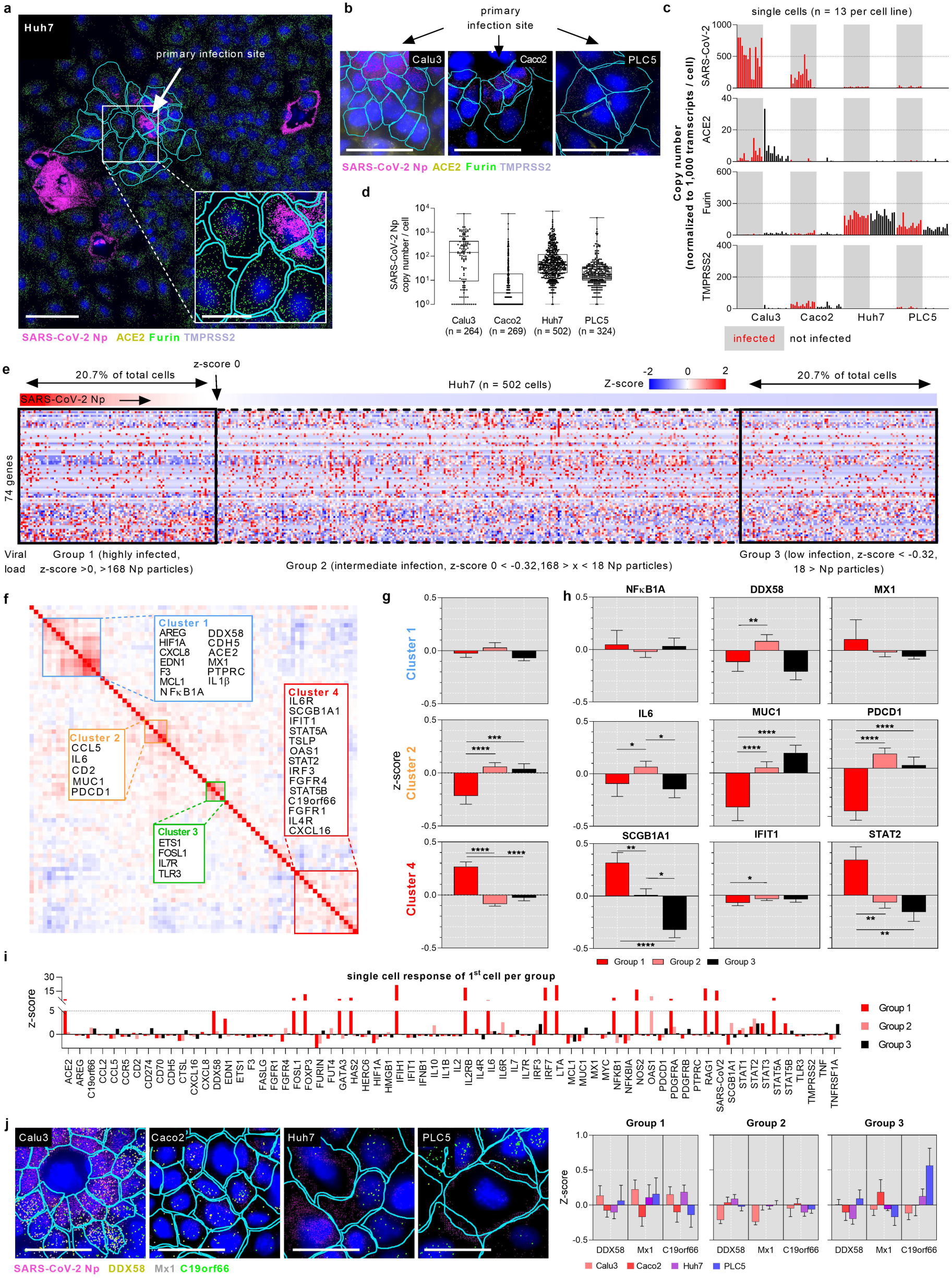
Infectivity of different cell lines with SARS-CoV-2 and spatial transcriptomics of key antiviral response genes. **a**, Huh7 cells infected with SARS-CoV-2. The primary infected cell (arrow) is surrounded by several secondary infected cells. Cell borders are outlined in blue. Transcripts of SARS-CoV-2 Np (magenta) and key SARS-CoV-2 entry factors *ACE2 (*yellow), *FURIN* (green) and *TMPRSS2* (gray) are highlighted. Scale bar, 100 μm; inlay scale bar 50 μm. **b**, *ACE2, FURIN* and *TMPRSS2* RNA species highlighted in Caco2, Calu3 and PLC5 cells. Scale bars, 50 μm. **c**, RNA species of SARS-CoV-2, *ACE2, FURIN* and *TMPRSS2* in infected and mock-infected Calu3, Caco2, Huh7 and PLC5 cells (n = 13 individual cells per cell line and sample). **d**, SARS-CoV-2 Np copy number per cell in infected Calu3 (n = 264), Caco2 (n = 269), Huh7 (n = 502) and PLC5 (n = 324) cells. Box plots comprise data from two independent biological replicates (n = 2) and indicate the overall mean ± SD. **e**, Heatmap of infected Huh7 cells (n = 502) ordered by decreasing number of detected SARS-CoV-2 Np RNAs per cell and hierarchical clustering of 74 different RNA species. Cells with a z-score of >0 (corresponding to >168 SARS-CoV-2 Np particles) were defined as highly infected and were classified as group 1, representing 20.7 % of the total evaluated cells overall. These were hypothesized to mainly contain primary infected cells. The 20.7 % of cells at the lower end of the heatmap were therefore inversely considered as slightly or not infected (z-score < -0.32, < 18 Np particles) and classified as group 3. Cells containing between 168 and 18 SARS-CoV-2 Np particles (z-score 0 > - 0.32) constituted group 2 and were considered as moderately or secondary infected. **f**, Matrix of all pairwise correlation coefficients of the expression of genes shown in e. Correlated genes are clustered by color as cluster 1 (blue), cluster 2 (orange), cluster 3 (green) and cluster 4 (red). **g**, Overall expression of genes per cluster sub-grouped in group 1, group 2 and group 3. Genes in cluster 2 and 4 were inversely deregulated in group 1 when compared to both groups 2 and 3, indicating expressional regulation depending on the infection rate. Data shown as mean ± standard error of the mean (SEM). **h**, Expression levels of selected genes per cluster comparing cells in group 1 – 3. Data shown as mean ± SEM. Significance levels in g. and h. were calculated by nonparametric Kruskal-Wallis test with Dunn’s test for multiple comparison. *p<0.05, **p<0.01, ***p<0.001 and ****p<0.0001. **i**, Single cell response of multiple genes of one cell per group. **j**, Antiviral response in different cell types. Left panel, close-up images of cells vicinal to the primary infection site (SARS-CoV-2 Np transcripts highlighted in magenta). Transcripts of *DDX58* (yellow), *MX1* (gray) and *C19orf66* (green) are highlighted. Scale bars, 50 μm. Right panel, comparison of expression patterns of *DDX58, MX1* and *C19orf66* across four cell lines subdivided by infection state as indicated by group 1-3. Data shown as mean ± SEM.

However, *ACE2* was the most prominent in Calu3 cells that also showed the highest amount of SARS-CoV-2 particles (normalized to total transcripts per cell). While *FURIN* was expressed at moderate levels across both hepatocellular cell lines and was not affected by the infection state, *TMPRSS2* was scarcely expressed in PLC5 and was absent in Huh7 cells. Interestingly, expression of both *FURIN* and *TMPRSS2* was slightly higher in non-infected Calu3 cells than in infected cells, indicating potential repression upon SARS-CoV-2 infection. However, although virus-mediated down-regulation of entry factors has been previously reported^31^, we observed no significant deregulation in our experiments. To assess cell-type dependent differences in overall infectivity, we compared total SARS-CoV-2 Np copy numbers per cell within the entire sample population. Notably, Calu3 cells showed the highest rates of viral particles per single cell (369.7 ± 664 particles, mean ± S.D., n = 264), while Caco2 cells seemed the least infected (65.95 ± 380.6, n = 269, **Fig. 3d**). Interestingly, Huh7 cells contained a substantially higher number of viral Np RNAs (168.9 ± 465.6, n = 502) per infected cell than PLC5 cells (61.8 ± 255.9, n = 324).

Next, we investigated the transcriptional responses to SARS-CoV-2 infections using a set of 81 genes in a single MC experiment and evaluated these relative to the viral load to assess potential virus-induced deregulation. A representative result from Huh7 cells (n = 502 single cells) is shown in **Fig. 3e**. Genes expressed in less than 10 % of cells (if 0 transcripts detected) were excluded ensuring deliberate analysis of antiviral genes relevant to the solid majority of cells (i.e. 74 genes were evaluated in Huh7 cells). Counts were normalized to the total transcripts per cell (with exception of SARS-CoV-2 Np transcripts to prevent skewing of data in highly infected cells), and scaled within the individual gene. Using a semi-supervised clustering approach, the cells were ordered by decreasing z-scores of viral transcripts (top lane) and clustered using a one-minus-Pearson correlation. We classified cells with a SARS-CoV-2 z-score value of 0 (equivalent to 168 viral particles per cell in Huh7 cells, termed group 1) as highly or primary infected, which accounted to 20.7 % of cells of the total population. The 20.7 % of cells at the very low end were regarded as lowly or not SARS-CoV-2 replicating cells (z-score of -0.32, < 18 viral Np particles, group 3) and were considered as bystander cells. Consequently, cells containing between 168 and 18 viral particles (z-score 0 > -0.32, group 2) were considered as moderate or secondary infections. Remarkably, the increased rate of highly infected cells of 37.5 % in Calu3 cells (> 392 viral Np particles, group 1) compared to 14.5 % in Caco2 (> 65 viral Np particles) and 11.5 % in PLC5 cells (> 62 viral particles) using identical MOI for infection complements the spread of SARS-CoV-2 via airborne transmission, as previously reported^32^. However, it was baffling that we observed no specific threshold of SARS-CoV-2 Np particles at which cells clearly react to the infection with a pronounced primary or secondary antiviral response.

Tempted by this observation, we went on to dissect the expression patterns at various stages comparing population-wide data in bulk analysis down to individual cells. We started by performing a pairwise correlation analysis and found four clusters of substantially co-regulated genes (minimum of four genes per cluster, **Fig. 3f**). In addition to genes coding for transcription factors and proteins regulating survival and apoptosis, cluster 1 comprises genes characteristic for SARS-CoV-2 infection such as *ACE2, DDX58* and *MX1*, the latter two exemplifying ongoing antiviral defence^30,33^. While cluster 2 and 3 reveal a diverse set of immune regulatory chemo- and cytokines (*CCL5* and *IL6*) as well as receptors (*IL7R, TLR3*), cluster 4 clearly points to the role of JAK/STAT signaling in host immune responses against viruses. With numerous combinations of STAT/STAT transcription factor complexes at the dispense of innate immunity, this highly complex network regulates the expression of hundreds of interferon stimulated genes (ISGs) that are imperative to curb, and ultimately clear viral infections^18,33,34^. For example, sensing of double stranded RNA by RIG-1 (encoded by *DDX58)* and endosomal TLRs, activates IRF3, IRF7 and NFκB, which results in the expression of type-I interferons (IFNα). These trigger the phosphorylation and therefore activation of a diverse set of STAT family members that promote the transcription of effector ISGs such as *MX1* or *OAS1* in an auto- and paracrine fashion. This contributes to priming of neighboring cells towards a collective antiviral response^33^. We assessed SARS-CoV-2-mediated deregulation of these clusters and implicated genes by comparing the previously defined groups (**Fig. 3g**). Genes in cluster 1 remained widely unaffected by varying infection rates while cluster 2 and 4 were significantly deregulated in highly infected cells, yet diametrically opposite.

We next tested whether differences observed on the level of gene clusters are also represented in the expression of selected individual genes (**Fig. 3h**). Surprisingly, *DDX58* expression declined in group 1 compared to group 2 which may indicate virus-induced downregulation that was not observed for *MX1*. Furthermore, *MUC1* and *PDCD1*, genes encoding cell surface proteins crucial in mediating inflammatory processes^35,36^, were significantly downregulated in group 1 suggesting interference of SARS-CoV-2 with diverse immune-regulatory processes beyond antiviral signaling. *SCGB1A1*, a component of cluster 4 that codes for a pulmonary surfactant protein, was recently suggested as a novel target for COVID-19 treatment due to its anti-inflammatory function^37^. Elevated levels of *SCGB1A1* observed in Huh7 cells that were lacking in Calu3 and Caco2 cells may thus support previous findings on the severity of organ-specific manifestations of COVID-19, which reported rather low levels of SARS-CoV-2-mediated liver injury in non-hospitalized patients^38,39^, potentially due to mitigation of hepatic inflammation by *SCGB1A1*. Moreover, the strong upregulation of *STAT2* expression and the downregulation of the ISG *IFIT1* in highly infected cells strongly supports viral interference at the level of nuclear translocation rather than transcription^40^. However, *IFITs* are also directly induced by IRF3 ensuring effective antiviral activity already during early stages of infection and are thus also expressed independently of type-I IFN signaling^41^.

Previous analysis demonstrated the application of MC in semi-bulk analysis at both the cell and target level. To further exploit the potential of MC, we set out to analyze expression levels of various genes on a single cell level (**Fig. 3i**). Notably, several genes were differently regulated on a single cell basis when compared to bulk analysis such as *DDX58, SCGB1A1* and *STAT2*, further emphasizing the role of single cell analysis in infection studies.

Remarkably, MC allows the simultaneous analysis of multiple samples at once, which is pertinent for directly comparing antiviral responses across various cell lines. Targeting *DDX58, MX1* and *C19orf66* concurrently in all four cell lines (representative images in **Fig. 3j**, left panel) grouped by the infection state, we found highly dispersing expression signatures across cell types and groups indicating differential regulation (**Fig. 3j**, right panel). Overall, counts in group 2 seemed to be the least deregulated with exception of downregulated *DDX58* and *MX1* in Calu3.

### Profiling of spatial response dynamics in primary INFS

We next used MC to investigate the viral propagation network including the coherent spatio-transcriptional response emanating from the primary infected cell. The spread of viral particles occurs either by diffusion, surface retention and cell-cell adhesion or cell-cell adhesion and polarization^42^ while viruses such as vaccinia virus induce virion repulsion to accelerate virus spread^43^. In an attempt to investigate the dissemination of SARS-CoV-2 particles within a single INFS, we evaluated viral particles including the coherent transcriptional antiviral responses in neighboring cells by dividing primary INFS in Calu3 cells into four regions of interest (ROI) separated by a distance of about one cell diameter (15 μm intervals, **Fig. 4a**). Here, two juxtaposed INFS are depicted showing the SARS-CoV-2 Np transcripts (magenta) as well as transcripts of *DDX58* (yellow), *MX1* (gray) and *C19orf66* (green). Note that, although INFS 2 contains a similar level of SARS-CoV-2 Np particles in ROI 1, the antiviral response in neighboring cells adjacent to the primary infected cell (ROI 0) is substantially less pronounced than in INFS 1. Based on this observation, we hypothesized that INFS 2 emerged at a later time point and hence is in an earlier state of viral replication. This may be due to a postponed replication of virus particles by a lower number of SARS-CoV-2 virions during initial infection, or due to INFS 2 starting as a secondary infection conveyed by diffusing particles emerging from INFS 1. To further explore this hypothesis, we compared the expression levels of target genes between the INFS attributed to their spatial distribution within the ROIs (**Fig. 4b**). Initially, endosomal entry of SARS-CoV-2 activates pattern recognition receptors (PRRs) such as TLR3 which leads to phosphorylation of IRF3 and IRF7 that activate expression of IFNα and, via JAK/STAT signaling, the expression of ISGs^33^. *TLR3* was strongly upregulated in ROI 2 in INFS 1 and moderately in INFS 2. Expression of *IRF3* and *IRF7* in ROI 2 was weaker in INFS 1 than in INFS 2 suggesting a delayed or even curbed IFNα response to emerging SARS-CoV-2 virions which indicates viral propagation to ROI 2^14,44^. *STATs* were highly expressed in ROI 1 in both INFS but only INFS 1 showed consistent elevation extended to ROI 2. Regarding *IFIH1* and *IFIT1*, potent ISGs involved in early stage viral inhibition, we found upregulation in ROI 2 only in INFS 1. Overall, the findings that upregulation of antiviral response genes in INFS 1 progressed to ROI 2 while remaining restricted to ROI 1 in INFS 2 supports the hypothesis of differently advanced infection events and advocates the necessity of highly-resolved multiplexed transcriptomics in studies on viral infection and replication.

**Fig. 4.**
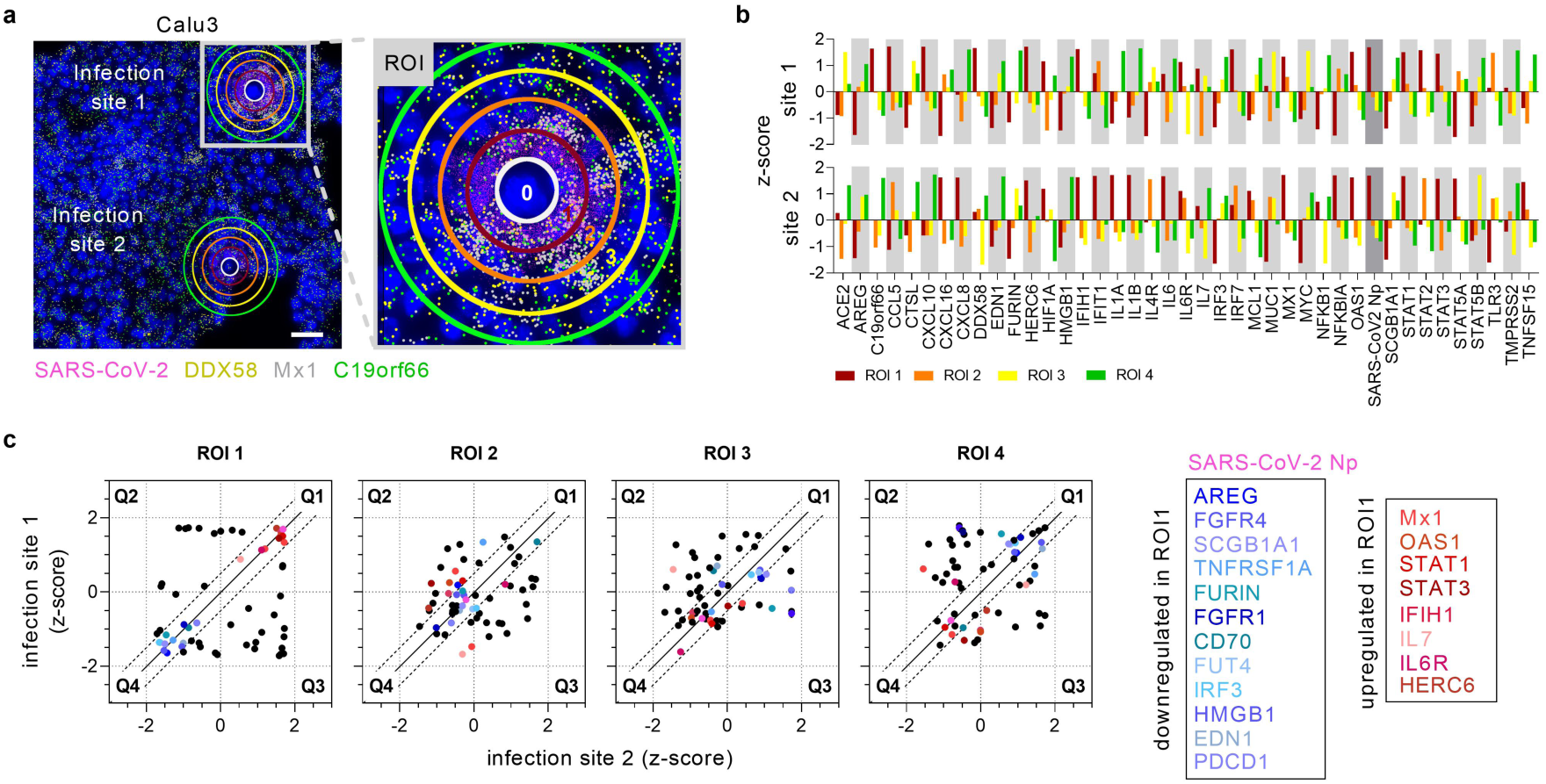
Spatially resolved, multiplexed profiling of response dynamics in primary infection sites (INFS). **a**, Representative image of two SARS-CoV-2 INFS in Calu3 cells with regions of interest marked by the primary infected cell (ROI 0), and outbound propagation of viral particles across ROI 1 – 4 including the coherent expression of antiviral genes. Highlighted RNA species comprise SARS-CoV-2 Np (magenta), *DDX58* (yellow), *MX1* (gray) and *C19orf66* (green). Scale bar, 50 μm. INFS 1 is enlarged for better visualization. **b**, Expression of multiple genes per ROI comparing INFS 1 and 2 (z-score calculated per infection site individually). **c**, Response trajectory of genes deregulated in both INFS across ROI 1 to ROI 4. Genes in Q1 and Q4 within 0.5 deviation (dashed line) from x = y (solid line) were considered for further analysis. Genes collectively upregulated in ROI 1 (Q1, red colors) tend to normalize their expression in ROI 2 and become downregulated in ROI 3, which seems to be fostered in ROI 4. The inverse pattern was observed for genes downregulated in ROI 1 (Q4, blue colors). Genes marked by the coherent colors are stated on the right side of panel c.

Lastly, to identify whether genes are mutually regulated in both INFS despite the deviating stage in viral propagation, we correlated the expression of all target genes between the INFS and traced their expression signatures across all four ROIs (**Fig. 4c**). We selected genes for tracing which were located in ROI 1 either in quadrant Q1 (upregulated, red colors) or Q4 (downregulated, blue colors) and were correlated with a maximum deviation of a z-score of 0.5 (dashed line) to the x = y axis (solid line). Remarkably, genes upregulated in ROI 1 tend to normalize their expression in ROI 2 and seem to become downregulated in ROI 3, a trend that was enhanced in ROI 4. This pattern was inversely true for genes downregulated in ROI 1. Similar to Huh7 cells, major upregulated genes in Calu3 cells within ROI 1 contained antiviral effector genes (*MX1, OAS1*) and members of the JAK/STAT signaling pathways (*STAT1, STAT3*) including their downstream-activated ISGs (*IFIH1*). Since we observed viral Np particles spreading in a rather circular pattern to the primary infected cell, we suggest that under the applied cell culture conditions, SARS-CoV-2 spreads primarily by conventional cell-to-cell adhesion through iterative rounds of infection, replication and release.

Together, these findings emphasize the role of STAT-pathway mediated antiviral defense and highlight the vast advantages of MC as a spatial, highly-resolved and multiplexed RNA profiling technique that allows the simultaneous study of early and late stage responses in various samples and target sites.

## Discussion

Spatial transcriptomic technologies are highly relevant for understanding biological behavior and architecture of cells and tissues as they allow the elucidation of location-dependent gene expression changes of multiple genes within cell communities or tissues. We developed a new technology, called MC, to complement the field of spatial transcriptomics with an approach that can be rapidly tailored to specific research needs and is particularly targeted at detecting and quantifying rare transcripts that are often missed in other applications^5,13,45^, necessitating both surpassing sensitivity and specificity. MC plugs into the standard smFISH workflow as a multiplex technology, theoretically providing the entirety of its advantages.

To verify the specificity of MC, we measured false positives (codes not in use) and misidentification rates (codes in use but no transcripts are expected) that represent meaningful specificity criteria and are subject to the whole MC process of hybridization, colorization, coding and de- and recoding. Considering the rate of misidentification, probe sets designed for human and mouse RNA transcripts applied to both human HeLa and mouse NIH-3T3 cells revealed a very low number of decoded signals (∼0.1 to 0.5 %) in non-target cells, thereby confirming the low rate of false positives (codes not in use). Remarkably, the number of false positives is 500 to 1000-fold lower than true positives as indicated by codes in use. Together, this ensures that a positive signal within the cell truly represents a specific transcript of the decoded gene. Concerning sensitivity, *Miedema et al*. reported smFISH to be one of the most sensitive spatial RNA detection methods and highly suitable for low abundance genes^46^, considering it to be the gold standard of spatial *in-situ* methods. Therefore, we compared MC with smFISH in their ability to detect two transcripts (*APC* and *CENPF*) against the background of multiple others and found their sensitivity to be indeed indistinguishable. This is supported by NGS data from RNAseq analysis that revealed an FPKM value of 7.6 for the *APC* gene and 37.1 for *CENPF*^47^. Since the coefficient of variation of *APC* detection was similarly low (∼0.12) compared to the more abundant *CENPF* transcript (∼0.11, **Fig. 1f**) in our experiments, despite being performed on several MC instruments and by different users, the quantification of transcripts of much lower abundance was considered to work equally well using smFISH and MC.

To demonstrate the vast applicability of MC, we localized both abundant and rare transcripts mapped at their subcellular resolution, including SARS-CoV-2 Np transcripts and dissected coherent antiviral responses of multiple genes in four different cell lines at once, which allowed evaluating response trajectories and demonstrating viral induced regulation of pathways affected by SARS-CoV-2. Remarkably, MC allows the analysis of both infected and non-infected cells directly within the same culture allowing the tracing of effects induced by paracrine signaling. Our study corroborates several aspects of viral interference with the host defense mechanism reported at the leading edge of SARS-CoV-2 studies. In localization studies, MC detects SARS-CoV-2 Np transcripts adjacent to the nucleus, suggesting ongoing translation of viral proteins in preparation for exocytosis^28^. The deviating nature of COVID-19 manifestations in multiple organs necessitates investigations in several cell types in direct side-by-side comparison^38,48^. MC facilitates such analysis and indeed revealed deviations in infectivity rates in cell lines of different origin.

These were complemented by detecting and allocating transcript counts of key entry factors such as *ACE2* that were so far reported as low or undetectable^14^, further illustrating the sensitivity of MC for rare targets. We found moderate to high infection rates of SARS-CoV-2 depending on the cell type, which allowed investigating viral propagation at various stages. In line with previous studies, we reported several high-interest targets to be deregulated in a sub-population of highly infected cells, most importantly members of type-I IFN stimulated pathways alongside downstream-activated ISGs^14,49,50^. Especially in the context of dynamic cell-to-cell or cell-to-pathogen mediated signaling, MC allows the mapping of both response propagation and trajectory, hence pinpointing at induction the antiviral state in bystander cells induced by paracrine signaling. Studying cells along this very edge of the infection gradient may help to better understand the interaction between effective viral clearance and immune escape mechanisms observed widely in COVID-19^51^.

MC provides several additional advantages, as it, for example, holds the potential to analyze multiple samples at once allowing direct side-by-side comparison; or appending immunostaining to exactly correlate transcriptomics data to expression patterns of related proteins. A pivotal asset of MC is the flexibility in target probe design that allows to quickly readjust working hypotheses, thus expediting day-to-day research. In practical terms, MC needs a single hybridization step of gene-specific probes with short steps of de- and re-colorization, followed by automated imaging. Although the MC experiments reported in this study were limited to a number of about 80 target genes related to SARS-CoV-2 induced cellular responses, a broad spectrum of future applications is envisaged.

Overall, MC is very flexible and highly suitable for multiple qualitative and quantitative applications of spatial transcriptomics approaches; particularly in studies targeting rare transcripts by providing utmost specificity, sensitivity, and reproducibility at high resolution (see **Fig. 1e to g**). By multiplexed detection of these transcripts at high resolution, MC promotes our ability to study subtle yet sophisticated cellular reactions especially in dynamic biological and medical processes.

## Online Methods

### Cell culture

HeLa (DSMZ, Cat. No. ACC 57) and NIH-3T3 cells (DSMZ, Cat. No. ACC59) were cultured in DMEM supplemented with 10 % FCS, 1 % penicillin-streptomycin (P/S, 10,000 U/mL), 200 mM L-Glutamine and 1 % MEM Non-Essential Amino Acids Solution (100X, all media and components from Gibco, ThermoFisher Scientific). African green monkey kidney epithelial Vero E6 cells (Biomedica, VC-FTV6) were maintained in MEM (Gibco), 5 % FCS and 1 % P/S. The following human cell lines were purchased from the Center for Medical Research (ZMF, Graz, Austria) and verified by short tandem repeat (STR) DNA profiling (ZMF). Human lung adenocarcinoma cell line Calu3 and human colon adenocarcinoma cell line Caco2 were maintained in MEM with 10 % FCS, 1 % P/S and 200 mM L-Glutamine. Human hepatocellular carcinoma cell line Huh7 and hepatoma PLC5 cells were maintained in DMEM with 10 % FCS and 1 % P/S. All cells were maintained at 37°C and 5 % CO_2_.

### Probe design

Multiple probes per targeted transcript were designed using an automated probe-designer software from Resolve BioSciences. Briefly, the probe-design was performed at the gene-level using all full-length protein-coding transcript sequences from the ENSEMBL database tagged as ‘basic’^52,53^. To speed up the process, the calculation of computationally expensive off-target searches, highly repetitive regions were filtered via the abundance of k-mers in the background transcriptome using Jellyfish^54^. A probe candidate was generated by extending a seed sequence until a certain target stability was reached. A set of rules was applied to discard sequences containing homopolymers, di- and trinucleotide repeats and strong compositional bias (e.g., lack of one base).

After these fast screens, every kept probe candidate was mapped to the background transcriptome using ThermonucleotideBLAST^55^ and probes with stable off-target hits were discarded. Specific probes were then scored based on the number of on-target matches (isoforms), which were weighted by their associated APPRIS level^56^, favoring principal isoforms over others. A bonus was added if the binding-site was inside the protein-coding region. From the pool of accepted probes, the final set was composed by picking the highest scoring probes.

### Sample preparation

Sample preparation of cultured cells on the slides was performed as described in the handbook *Cell preparation for Molecular Cartography*. Briefly, approximately 1 × 10^4^ cells (defined per cell line) were seeded per well on MC slides and grown to 30-50 % confluence. The cells were grown for 1 or 2 days at 37 °C in the appropriate medium. After the medium was aspirated, cells were washed and fixed using methanol at -20 °C for 10 min. Slides were dried and stored at -80 °C.

### Hybridization of transcript specific probes

All steps were performed exactly according to the protocol of the *Molecular Cartography Hybridization (Cells)* Kit. Briefly, the cells were pretreated with 70 % ethanol followed by solution BSC1, and wash buffer 1. Thereafter, gene-specific probes were hybridized for 15-18 h at 37 °C in pre-warmed hybridization solution. Subsequently, the cells were washed twice using wash buffer 2 for 10 min each before the temporary addition of wash buffer 1 before color development. The storage buffer may be applied to store the slide at 4° C for up to two weeks. All buffers, solutions, and reaction conditions are described in the *MC Hybridization (Cells)* protocol.

### Color development and Decolorization

The color code of the hybridized probes develops over eight cycles to efficiently distinguish the individual transcripts. Therefore, the following protocol was repeated eight times. For round 1, color-solution 1 was prepared to contain the reagent CR1. After incubating the hybridized cells with color-solution 1 for 45 min at 25° C, a washing step was performed to eliminate excess reagent. Thereafter, color-solution 2 was applied containing the reagents CD1 and CD2 and incubated for 45 min at 25° C. Subsequently, cells were washed three times followed by the addition of imaging buffer and microscopic imaging. After the 1^st^ round of imaging, a second round is initiated using reagent CR2. Before the second round, a decolorization of 1^st^ round color is performed. For decolorization, the cells were washed five times using pre-warmed wash buffer 1 for 6 min at 41° C before proceeding with the next colorization round using a freshly prepared color solution 1 containing the CR2 reagent. Washing steps, the incubation with color-solution 2, and imaging is performed as described above. For the following rounds, the reagents CR3 to CR8 are used. All steps, buffer ingredients, colorization, and decolorization solutions are stated in the protocol provided by Resolve BioSciences.

### MC Imaging

The imaging was done by a customized Zeiss Celldiscoverer 7 (CD7) device, using the 50x Plan Apochromat water immersion objective with an NA of 1.2 and the 0.5x magnification changer, resulting in a 25x final magnification. Standard CD7 LED-light source, emission filters, and dichroic mirrors were used together with customized emission filters optimized for detecting the specific signals. Excitation times per image were 1000 ms for each channel (DAPI was 20 ms). A z-stack was taken at each region with a distance of 300 nm per z-slice. The custom CD7 CMOS camera (Zeiss Axiocam Mono 712, 3.45 μm pixel size) was used resulting in an image pixel edge length of 138 nm. The total size of an image was 295.9 μm x 295.9 μm, defining the size of one imaging region.

For each region to be analyzed, a z-stack for each of the two fluorescent colors was imaged in every imaging round. A total of 8 imaging rounds was performed for each position, resulting in 16 z-stacks per region. The fully automated imaging process (including water immersion generation and precise relocation of regions to image in all three dimensions) was realized by a custom Python script using the scripting API of the Zeiss ZEN software (open application development).

### Primary analysis of raw data

The primary data analysis was done in several steps, which are detailed below: Preprocessing: All images were corrected for background fluorescence. Based on the raw data image, the 20 % darkest local pixel values and positions were determined and copied to a new empty image (background image) having the same size as the image to be corrected. The remaining 80 % of pixels of the background image were generated based upon the surrounding existing pixel values using a distance weighted average value. Finally, the background-corrected image (bc-image) was created by subtracting the background image values from the raw data image values. The bc-image was used for all subsequent analysis steps.

### Extraction of features from the bc-images

In a first step, a target value for the allowed number of maxima was calculated based on the area of the slice in μm^2^ multiplied by an empirically optimized factor (0.5x). The resulting target value was used to adapt the threshold for the algorithm iteratively searching local 2D-maxima. The threshold leading to the closest number of maxima equal to or smaller than the target value was used for further steps and the respective maxima were stored. This procedure was done for every image slice independently. Maxima that did not have a neighboring maximum in an adjacent slice (called z-group) within a radius of one pixel were excluded. For the resulting list of maxima, the absolute brightness (Babs), the local background (Bback), and the average brightness of the pixels surrounding the local maximum (Bperi) were measured and stored. The resulting maxima list was further filtered in an iterative loop by adjusting the allowed thresholds for (Babs-Bback) and (Bperi-Bback) to reach a feature target value based upon the total volume of the 3D-image. Only maxima still in a z-group with a size of at least 2 were passing the filter step. Each z-group was counted as one hit. The members of the z-groups with the highest absolute brightness were used as features and written to a file. These features resemble 3D point clouds.

### Iterative closest point algorithm for determination of transformation matrices

To align the raw data images from different imaging rounds, these images had to be corrected for the 6 degrees of freedom in 3D-space. To do so, the extracted feature point clouds were used to find the transformation matrices. For this purpose, an iterative closest point cloud algorithm was used to minimize the error between the point clouds. Prior to the alignment the feature point clouds of the different colors in one round were merged based upon the assumption to be aligned already. The point clouds of each round were aligned to the point cloud of round one (reference point cloud). The sequence of alignments was always from round 2 to round 8. The points from round 2 to 4 were added to the reference point cloud, when no point of the current reference point cloud was in vicinity after transformation. The resulting transformation matrices were stored for downstream processes. Based upon these transformation matrices a transformation of the corresponding images was performed.

### Pixel evaluation

The aligned images were used to create a profile for each pixel consisting of 16 intensity-values, one from each of the 16 images (two colors, eight rounds). Pixel profiles with low variance were removed and the remaining profiles were compared to each profile of the encoding scheme used in the experiment. Profiles were then further filtered based upon similarity between best and second-best match, as well as by their ratio of summed signals that are contributing to the matched profile (constructive signal) to summed signals that were not expected (noise signal) based on the encoding schema. The match with the highest score (constructive signal minus noise signal) was assigned as an ID to the pixel.

### Filtering and evaluating pixel groups

Pixels with neighbors having the same ID were grouped. The pixel groups were filtered by group size, number of direct adjacent pixels in the group, number of dimensions with size of at least two pixels. The local 3D-maxima of the groups were determined as potential final transcript locations. Maxima were excluded if there was no corresponding maximum in the raw data images. Remaining maxima were further evaluated by their fit to the corresponding code. The final list of maxima was written to the results file and considered to resemble transcripts of the corresponding gene. Not all possible valid words of the encoding schema are used in an experiment. The ratio of expected words (these are used in the experiment) and words matching to valid codes that were not used in the experiment were used to measure specificity and estimate false positives.

### Determining specificity

The specificity of the probe sets was evaluated by simultaneously hybridizing 16 human and 8 mouse genes to both cell types and testing each for positive and negative signals. Human Hela Cells and mouse NIH-3T3 cells were grown overnight as described above. Probe sets were hybridized and transcripts within the cells were detected according to the standard MC protocol. Counts for transcripts of human and mouse cells were determined individually. Three independent experiments were conducted (n = 3 biological replicates).

### Determining sensitivity

The sensitivity of MC was tested in comparison to smFISH. Human Hela cells were grown overnight as described above. For MC, probe sets for 19 human genes were hybridized to fixed HeLa cells and transcripts were detected using the standard MC protocol. For smFISH, we chose two probe sets per gene (each of which contains > 45 probes). Both probe sets are differentially labeled by different fluorophores. The probe sets were designed so that a probe with fluorophore A binds vicinal to a probe with fluorophore B. Both probe sets are hybridized simultaneously using the protocols and buffers offered by Biosearch technologies for smFISH experiments. A signal was evaluated as positive only if the signal was detectable by both fluorophores.

### Preparation of SARS-CoV-2 virus stock

SARS-CoV-2 (Human 2019-nCoV Isolate, Ref-SKU 026V-03883), as disclosed by the Center for Disease Control and Prevention, was obtained from Charité Universitätsmedizin Berlin. SARS-CoV-2 was grown in Vero E6 cells in EMEM supplemented with 10 % FBS and 1 % P/S, frozen and thawed once, centrifuged at 900 g and filtered through a 0.22 μm filter (ThermoFisher Scientific). Infection titers were determined by focus forming assay and TCID-50/mL calculated with Spearman-Kaber and Reed-Muench test.

Experiments of propagating and applying infectious SARS-CoV-2 were performed in the bio-safety-level 3 (BSL-3) facility at the Diagnostic & Research Center for Molecular BioMedicine at the Medical University of Graz, Austria, following institutional biosafety guidelines.

### Verification of SARS-CoV-2 infectivity

Infectivity of SARS-CoV-2 virus stocks was determined in VeroE6 cells by infecting cells grown to a confluence of 70-80 % at an MOI of 0.002 for 1 h, washed twice with PBS and incubated for 24 h at 37 °C, 5 % CO_2_. RNA was extracted from the supernatant using the QIAamp Viral RNA Mini Kit (Qiagen). Viral replication was quantified by RT-qPCR using the CDC 2019-Novel Coronavirus (2019-nCoV) Real-Time RT-PCR Diagnostic Panel (2019-nCoV_N1 Forward Primer 5’-GAC CCC AAA ATC AGC GAA AT-3’ and 2019-nCoV_N1 Reverse Primer 5’-TCT GGT TAC TGC CAG TTG AAT CTG-3’ with the 2019-nCoV_N1 Probe 5’-FAM-ACC CCG CAT TAC GTT TGG TGG ACC-BHQ1-3’, ^57^) on a Rotor-Gene (Qiagen) using the following protocol: hold 1: 30 min 50 °C, hold 2: 15 min 95 °C, cycling 3 s 95 °C, 30 s 55 °C for 45 cycles. A commercially available standard (Genomic RNA from 2019 Novel Coronavirus, ATCC, VR-1986D) was used to assess viral copy number. The infectivity of the virus stock was further verified by immunohistochemistry (IHC) using a primary rabbit monoclonal antibody against the SARS-CoV-2 (2019-nCoV) nucleocapsid protein (SinoBiological, 40143-R019) and commercial staining reagents (Agilent Technologies, K346430-2, K406189-2) following the manufacturer’s instructions.

### SARS-CoV-2 infection for MC

For SARS-CoV-2 infection experiments, 7.000 Huh7 cells, 15.000 PLC5 cells, 15.000 Caco2 cells and 20.000 Calu3 cells were seeded into 8x glass bottom slides (Ibidi) at a confluence of 30-50 % and infected with SARS-CoV-2 at an MOI of 0.4 (if applied to VeroE6 cells, titer maintained for comparison across cell lines) for 60 min at 37 °C at 5 % CO_2_. The cells were then washed two times with medium and incubated for 24 h at 5 % CO_2_ at 37 °C. Supernatant was collected to verify viral replication as stated above. The cells were fixed by incubation in ice-cold methanol for 10 min and ethanol for 1 min. The slides were stored at -80 °C until further use.

### Hierarchical clustering and covariation analysis

Expression data was normalized to the total counts per cell (with exception of SARS-CoV-2 Np transcripts to avoid skewing of data) or ROI (Fig. 4) and scaled using Excel (Microsoft).

Genes expressed in less than 10 % of cells (if 0 transcripts detected) were excluded to emphasize transcriptional signatures relevant to the solid majority of cells. This threshold can be adjusted to identify expression patterns of abundant or rare transcripts depending on the research question. Generation of heat maps and hierarchical clustering for all experiments was conducted using the open source platform Morpheus^58^. For heat maps, data was organized by decreasing number of SARS-CoV-2 z-scores and clustered by one-minus-Pearson correlation coefficient of the cell-to-cell variations of the detected copy numbers of each pair of RNA species. Hence, genes with strongly correlated expression patterns appear closer, while distantly correlated genes appear further apart. These distances were used for the construction of a pairwise correlation matrix. Groups with at least four genes with substantial covariations were selected for further analysis.

### Data analysis and visualization

Analysis of raw data was performed using excel (Microsoft, Washington, US) and GraphPad Prism (version 8.4.3,^59^) for statistical analysis and graphical representation. Data was tested for normal distribution using a Shapiro-Wilk normality test. Since no data set passed the normality test, we performed statistical analysis using non-parametric Spearman rank test (two-tailed) for correlation analysis and a Kruskal-Wallis test with Dunn’s multiple comparison test for data sets with group or target comparison as indicated in the figure legends. Statistical significance was considered at p < 0.05 (*), p < 0.01 (**), p < 0.001 (***) and p < 0.0001 (****). Final figures were prepared using GraphPad Prism.

### Reporting summary

Further information on research design is available in the Nature Research Reporting Summary linked to this article.

**Table 1:**
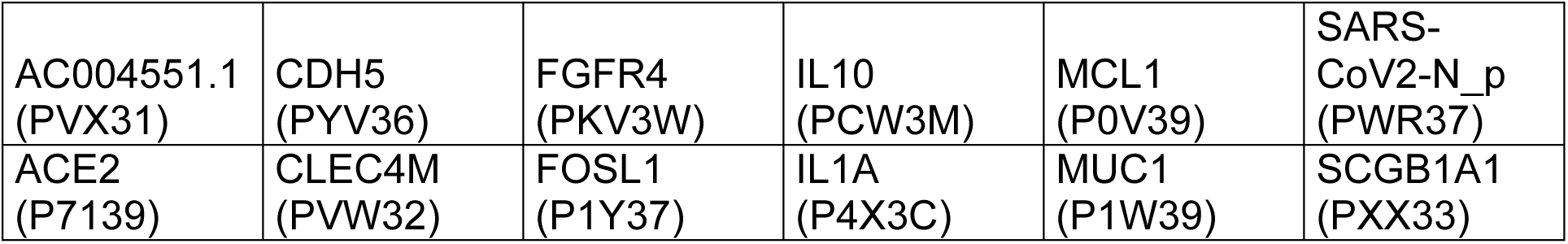
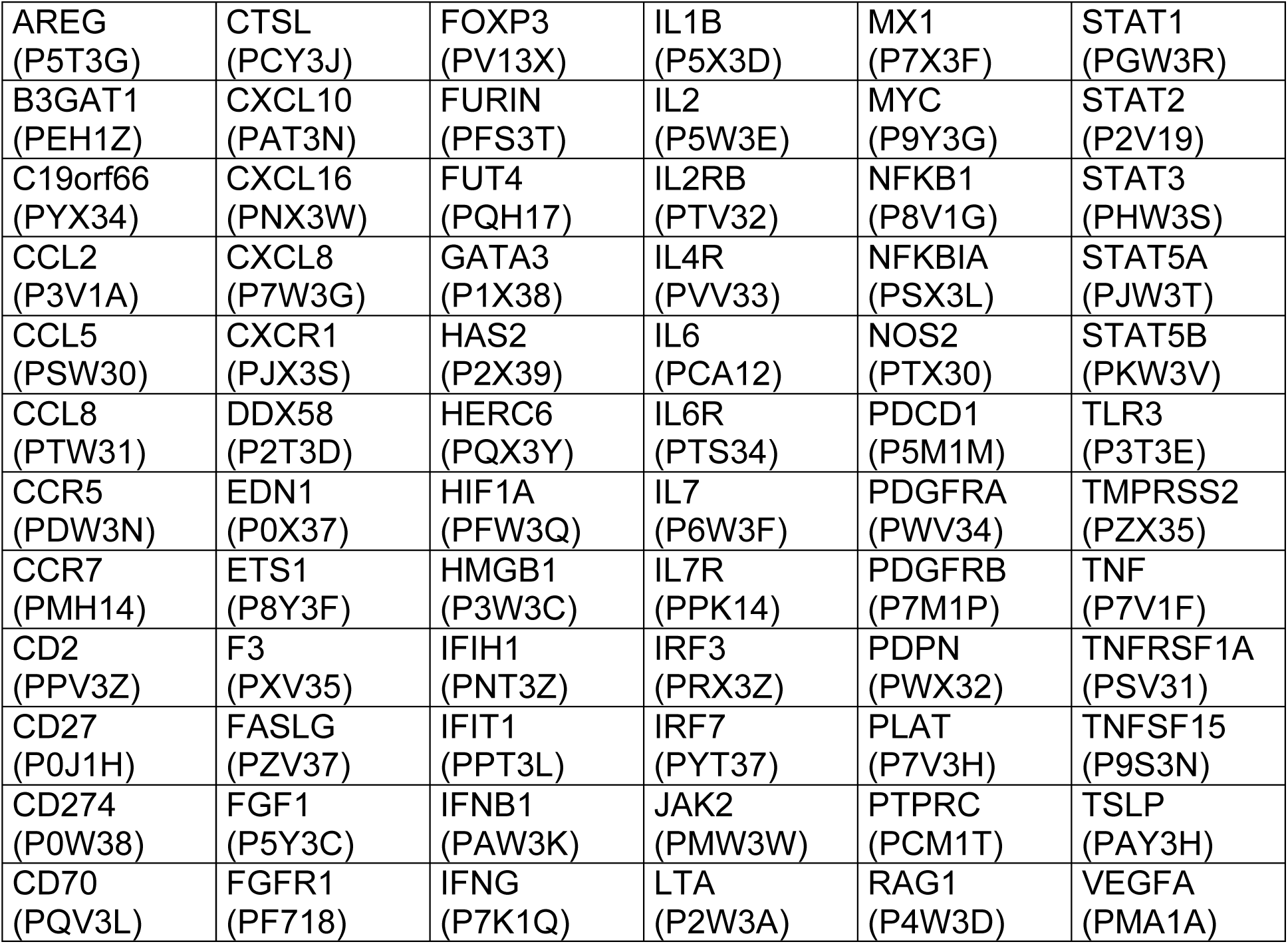
Probes (including catalogue numbers) used for infection experiment.

|  |  |  |  |  |  |
| --- | --- | --- | --- | --- | --- |
| AC004551.1<br>(PVX31) | CDH5<br>(PYV36) | FGFR4<br>(PKV3W) | IL10<br>(PCW3M) | MCL1<br>(P0V39) | SARS-CoV2-N_p<br>(PWR37) |
| ACE2<br>(P7139) | CLEC4M<br>(PVW32) | FOSL1<br>(P1Y37) | IL1A<br>(P4X3C) | MUC1<br>(P1W39) | SCGB1A1<br>(PXX33) |
| AREG<br>(P5T3G) | CTSL<br>(PCY3J) | FOXP3<br>(PV13X) | IL1B<br>(P5X3D) | MX1<br>(P7X3F) | STAT1<br>(PGW3R) |
| B3GAT1<br>(PEH1Z) | CXCL10<br>(PAT3N) | FURIN<br>(PFS3T) | IL2<br>(P5W3E) | MYC<br>(P9Y3G) | STAT2<br>(P2V19) |
| C19orf66<br>(PYX34) | CXCL16<br>(PNX3W) | FUT4<br>(PQH17) | IL2RB<br>(PTV32) | NFKB1<br>(P8V1G) | STAT3<br>(PHW3S) |
| CCL2<br>(P3V1A) | CXCL8<br>(P7W3G) | GATA3<br>(P1X38) | IL4R<br>(PVV33) | NFKBIA<br>(PSX3L) | STAT5A<br>(PJW3T) |
| CCL5<br>(PSW30) | CXCR1<br>(PJX3S) | HAS2<br>(P2X39) | IL6<br>(PCA12) | NOS2<br>(PTX30) | STAT5B<br>(PKW3V) |
| CCL8<br>(PTW31) | DDX58<br>(P2T3D) | HERC6<br>(PQX3Y) | IL6R<br>(PTS34) | PDCD1<br>(P5M1M) | TLR3<br>(P3T3E) |
| CCR5<br>(PDW3N) | EDN1<br>(P0X37) | HIF1A<br>(PFW3Q) | IL7<br>(P6W3F) | PDGFRA<br>(PWV34) | TMPRSS2<br>(PZX35) |
| CCR7<br>(PMH14) | ETS1<br>(P8Y3F) | HMGB1<br>(P3W3C) | IL7R<br>(PPK14) | PDGFRB<br>(P7M1P) | TNF<br>(P7V1F) |
| CD2<br>(PPV3Z) | F3<br>(PXV35) | IFIH1<br>(PNT3Z) | IRF3<br>(PRX3Z) | PDPN<br>(PWX32) | TNFRSF1A<br>(PSV31) |
| CD27<br>(P0J1H) | FASLG<br>(PZV37) | IFIT1<br>(PPT3L) | IRF7<br>(PYT37) | PLAT<br>(P7V3H) | TNFSF15<br>(P9S3N) |
| CD274<br>(P0W38) | FGF1<br>(P5Y3C) | IFNB1<br>(PAW3K) | JAK2<br>(PMW3W) | PTPRC<br>(PCM1T) | TSLP<br>(PAY3H) |
| CD70<br>(PQV3L) | FGFR1<br>(PF718) | IFNG<br>(P7K1Q) | LTA<br>(P2W3A) | RAG1<br>(P4W3D) | VEGFA<br>(PMA1A) |

**Table 2.** Probes used for HeLa cells (reproducibility and sensitivity)

|  |  |  |  |  |  |
| --- | --- | --- | --- | --- | --- |
| ACO2<br>(PJX1Q) | AURKB<br>(PQ61J) | BIRC5<br>(P1A1S) | CDC25C<br>(P4125) | DDX5<br>(PJ31G) | HPRT1<br>(P4X19) |
| AKT1<br>(PNT1X) | BCL2<br>(PT71M) | BRCA1<br>(PQ02S) | CDK1<br>(P6127) | E2F1<br>(PPA1D) | KRAS<br>(PCX1H) |
| AOX1<br>(PY02L) | BCR<br>(PZX13) | CCNE1<br>(P2123) | CENPF<br>(P8129) | EGFR<br>(PDX1J) | TP53<br>(PEX1K) |
| APC<br>(PJ71C) |  |  |  |  |  |

**Table 3:** Probes used for HeLa cells (specificity experiment)

|  |  |  |  |  |  |
| --- | --- | --- | --- | --- | --- |
| AURKA<br>(PWA1K) | CD70<br>(PQV3L) | DHFR<br>(PC22C) | IL4R<br>(PVV33) | MTIF3<br>(PPV1X) | WASF2<br>(PMT1W) |
| BUB1B<br>(PD32C) | CD83<br>(PKX3T) | GTSE1<br>(PC22C) | IL6 (PCA12) | PILRB<br>(PCV1K) |  |
| CCL2<br>(P3V1A) | CDKN1A<br>(P4A1W) | IFI44<br>(P4V1C) | MKI67<br>(P1Z14) | PLIN3<br>(P9C20) |  |

**Table 4:** Probes used for mouse NIH-3T3 cells

|  |  |  |  |
| --- | --- | --- | --- |
| AMIGO2<br>(P7Z2C) | GLRX3<br>(P8P1N) | IL18<br>(PPP55) | NEAT1<br>(PVP16) |
| CFD<br>(P8C1Z) | GSDMD<br>(PDD11) | MKI67<br>(P2Q3G) | PLIN3<br>(PTD2G) |

## Data availability

Data underlying the reported findings is available from the corresponding author on request.

## Acknowledgement

The authors are grateful to Penelope Kungl and Jason Gammack for editing the manuscript.

## Author contributions

S.G., C.K. and K.Z. wrote the manuscript. S.G. prepared the figures. C.K. and K.Z. designed and supervised the project. A.G., F.R., and C.K. developed MC technology. A.G., F.R., B.N. and C.K. performed gene selection for SARS-CoV-2 experiments. C.U. planned MC specificity and sensitivity experiments. C.U., C.F., A.M. performed MC specificity and sensitivity experiments. S.G., D.P., C.K, and K.Z. designed the experiment on SARS-CoV-2 infection. E.F.-H. and M.H. conducted SARS-CoV-2 propagation and infection. S.G., A.B., B.N. and D.P. conducted MC experiments on SARS-CoV-2 infection. C.K and S.G. performed data analysis. S.S. conducted bioinformatics analysis for the video.

All authors read and commented on the paper.

